# Supporting *Computational Apprenticeship* through educational and software infrastructure. A case study in a mathematical oncology research lab

**DOI:** 10.1101/835363

**Authors:** Aasakiran Madamanchi, Madison Thomas, Alejandra Magana, Randy Heiland, Paul Macklin

## Abstract

There is growing awareness of the need for mathematics and computing to quantitatively understand the complex dynamics and feedbacks in the life sciences. Although several institutions and research groups are conducting pioneering multidisciplinary research, communication and education across fields remains a bottleneck. The opportunity is ripe for using education research-supported mechanisms of cross-disciplinary training at the intersection of mathematics, computation and biology. This case study uses the computational apprenticeship theoretical framework to describe the efforts of a computational biology lab to rapidly prototype, test, and refine a mentorship infrastructure for undergraduate research experiences. We describe the challenges, benefits, and lessons learned, as well as the utility of the computational apprenticeship framework in supporting computational/math students learning and contributing to biology, and biologists in learning computational methods. We also explore implications for undergraduate classroom instruction, and cross-disciplinary scientific communication.

## 1 Introduction

Over the last several decades advances in experimental techniques have provided life scientists with increasing quantities of high dimensional, high-resolution datasets. Unfortunately these technological developments have not yet been matched by similar clinical advances [1]. In fact, U.S. life expectancy has stagnated and development costs for new drugs continue to rise [2, 3].

Over the last decade, a consensus has emerged among scientific thought leaders about the need for “convergence” or transdisciplinary integration of engineering and physical science approaches in the life sciences to generate meaningful biological insights from the growing abundance of biomedical data [4, 5]. In particular, computational modeling approaches, which include both mathematical modeling of complex cell and molecular systems and statistical modeling of large datasets, are a powerful vehicle for synthesizing disparate and sometimes conflicting data into an integrated biological understanding. Ideally, computational modeling approaches work recursively with experimental workflows; mathematical models quantitate and formalize the largely qualitative, observation-driven “mental models”, and the iterative comparison of model outputs to experimental data informs model refinement and also suggests new experimental directions [6–9].The use of mathematical models clarifies the biological conditions or parameters under which the “mental model” can explain the experimental and simulation data. Increasingly, statistical modeling approaches including machine learning and bioinformatics are used to complement mathematical modeling of cell and molecular biological systems [7]. Analysis of large clinical or experimental datasets can be used to inform parameterization of mathematical models, or to identify novel relationships between cell states and behaviors which can generate new hypotheses for mathematical modeling. Additionally, machine learning methods can be used for richer and more informative analysis of mathematical model outputs.

Despite the consensus around the need for greater integration of computational and experimental modeling approaches, there remains relatively limited adoption of these methods in the life sciences community. Furthermore, the research groups that employ computational modeling approaches largely work in isolation using their own data sources, building their own models, and performing their own analyses [7]. Biologists in experimental research groups face substantial structural, technical and educational barriers to learning to implement computational modeling approaches [10]. These challenges are compounded by resource limitations at emerging research institutions including minority serving institutions and primarily undergraduate institutions [11]. Expanding participation in computational modeling approaches requires adoption of innovative practices in cross-disciplinary scientific communication and training [10].

Similarly, the traditional educational divisions between engineering, computational, physical science and biological curricula have impeded communication between these silos and raised barriers both to the use of simulations by biologists and the effective understanding of biological needs and design of tools for biological applications by engineers [12, 13]. There has been an ongoing effort to incorporate quantitative or computational content into undergraduate biology courses [14–19]. Similarly, there are several reports of reforms to provide greater life sciences disciplinary content to engineering students [20–22]. Nevertheless, there is a need for theoretically sound, evidence-supported models for undergraduate classroom instruction and undergraduate research experiences to train new cohorts of biologists and engineers equipped to work at the intersection of computation and biology. Additionally, there is a need for collaboration with education researchers to develop and assess interdisciplinary efforts [23].

In this paper, a computational apprenticeship framework is used to present a case study of our computational biology research group’s experience in iteratively developing an educational infrastructure for undergraduate research experiences. We describe a mutually beneficial conversation and collaboration with computational education researchers to develop, assess, and continuously improve undergraduate research involvement in a research active computational laboratory.

We adopt *rapid prototyping* approaches from engineering to iteratively design, test, and refine the mentorship infrastructure: after implementing a current mentorship version in the lab, we evaluate strengths and weaknesses, identify concrete refinements to address weaknesses while building upon strengths, update the mentorship structure (with a new version number), and continue testing in the subsequent research term. This case study will present each mentorship version as we stepped through this iterative design process.

### 1.1 Computational Apprenticeship

*Computational Apprenticeship* is a newly proposed theoretical framework and a type of cognitive apprenticeship for computational disciplines [24]. Biology and engineering, like most other academic disciplines, are often taught through traditional instructional methods heavily featuring didactic lectures. In these disciplinary contexts, there is typically a heavy emphasis on the technical aspects of computational topics. However, education research suggests that a narrow focus on technical competency typically provides students with routine expertise, but lacks the adaptive expertise needed to solve computational problems in real world settings [24–26]. For example, students attempting to develop computational models of cellular and molecular biological systems report challenges with higher-order computational thinking skills such as abstraction and problem decomposition, rather than coding or mathematics [10, 27, 28].

Improving computational education requires greater consideration for helping students develop their ability to solve disciplinary problems with computation, rather than merely master the technical aspects of worked examples. Cognitive Apprenticeship is grounded in constructivist learning theories and draws on traditional apprenticeship training structures in the skilled trades. This model argues that students learn best through guided-experience on cognitive and metacognitive skills and processes specific to their discipline [29]. Collins et al. outlined Content, Method, Sequencing, and Sociology as the four critical dimensions of learning environments [30]. The Computational Apprenticeship framework adapts this model to computational domains and provides a theoretical basis for creating curated learning experiences with graduated challenges in terms of difficulty and diversity. The framework also provides direction for the use of emerging technological practices such as code commenting and Jupyter notebooks to deliver pedagogical scaffolding and other elements of the *sequencing* and *method* dimensions of learning [24].

### 1.2 Research group context

Macklin’s MathCancer Lab is a computational mathematical biology research group that develops theory- and data-driven computational model systems that can help understand and engineer the behavior of multicellular systems, especially in cancer and tissue engineering. Tackling these goals necessitates both multicellular systems biology and multicellular systems engineering perspectives [7, 31]. The development of next-generation cures requires a deeper understanding of the fundamental biology of multicellular systems [32]. Reductionist approaches— motivated in part by the earlier successes of germ theories for infectious diseases—attempt to cure diseases by identifying and repairing a single root cause (e.g., a single or small number of driver mutations, or an overactivated receptor pathway) in an isolated cell type [33, 34]. These approaches, however, neglect the complex interactions in the evolving multi-level networks of normal and diseased tissues [35]. Targeted interventions do not only affect just single cell types; biochemical and biophysical feedbacks—combined with intercellular heterogeneity and natural selection [32] and amplified by physical constraints [36]—can cause secondary effects such as therapeutic resistance (e.g., by selecting for resistant cancer clones), worsened drug delivery, and treatment toxicity [6, 32, 37]. Thus, next-generation therapies must not just treat single cell types, but rather steer the multicellular systems towards balance. This necessitates systems thinking that combines biological domain expertise with computational and mathematical tools designed for complex cell and molecular biological systems [6, 32], along with scientific computing infrastructures for large-scale investigations [6, 38].

To drive these systems approaches, the MathCancer Lab develops the technological core components and infrastructure of a computational model system that can be interrogated for multicellular systems biology and engineering from a multiscale mathematics perspective [32]. They developed an efficient open source framework to solve coupled reaction-diffusion partial differential equations in biochemical tissue environments [39], and an agent-based framework. (PhysiCell) for multiscale modeling that can include stochastic processes, systems of ordinary differential equations, and other mathematical “submodels” in thousands or millions of interacting discrete cell agents [40]. (See [41] for an overview of cell-based modeling methods in cancer.)

Together, these components form the backbone of a computational tissue model system for simulating and analyzing multicellular biological systems that combine discrete and continuum mathematics [32]. Typical applications include cancer immunology, synthetic multicellular systems, metabolic tumor-stroma crosstalk in heterogeneous cell populations, and infectious diseases. See some examples in Fig. 1.

**Figure 1.**
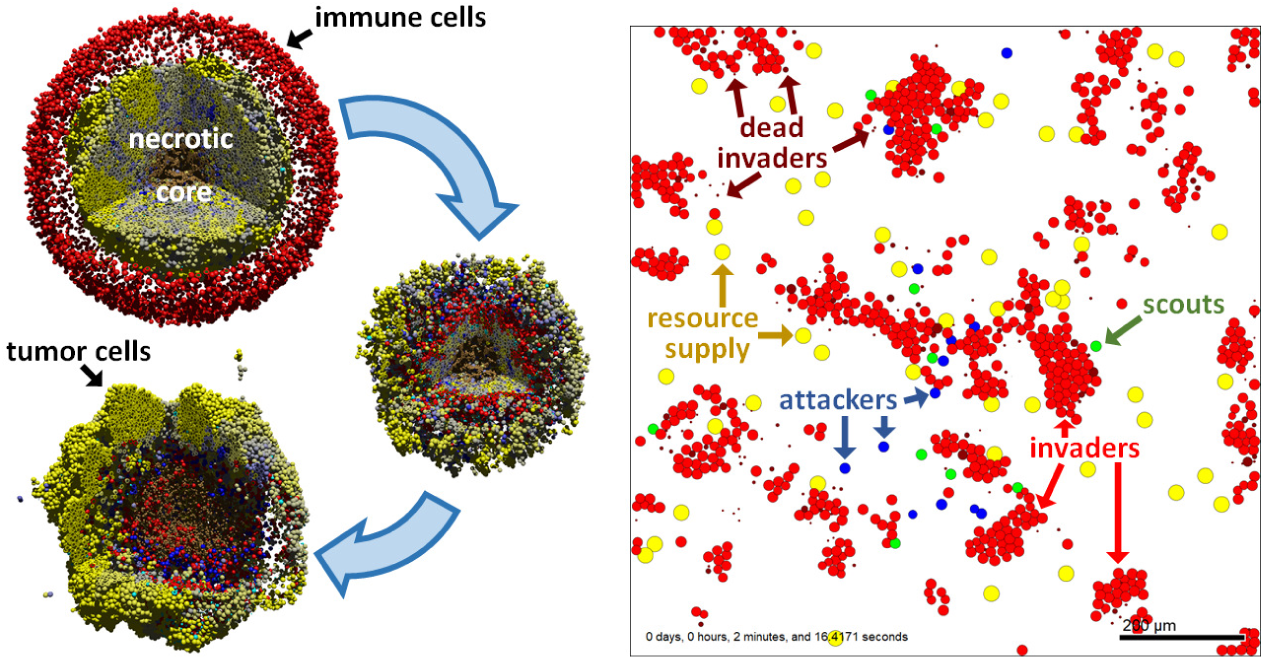
Sample PhysiCell models. <u>left</u>:Cancer Immunotherapy. See a full description and 3D visualization at [42] and a cloud-hosted 2D interactive version at [43]. Adapted under CC-BY license from [40]. right: Cell-cell communication by chemical diffusion. See a cloud-hosted interactive version and full description at [44].

Partnerships with the open source community have combined these modeling frameworks with large-scale investigations on supercomputers [6], machine learning approaches that accelerate the investigations and aid model interpretation [38], and new mathematical model components (e.g., Boolean signaling networks [45]). Efforts towards data standardization [7, 46] and a recent focus on creating open educational training materials and shared source code repositories seek to grow these computational projects from a single-lab effort to a community-driven ecosystem for mathematical biology, multicellular systems biology, and computational bioengineering [7, 47].

To advance towards these goals, the lab has recently focused on integrating undergraduate research, undergraduate education, and scholarly communication, drawing upon evidence-based practices from the computational apprenticeship framework. Undergraduate and graduate students, scientific staff, faculty, visiting researchers, and a network of multi-disciplinary collaborators work together to build computational resources, apply them to specific cancer and other biological problems, disseminate methods and results, and unite researchers for community-driven science. Here, we share our experiences in this iterative effort.

## 2 Case study: Undergraduate involvement in a computational oncology laboratory

The MathCancer Lab moved to Indiana University’s new Intelligent Systems Engineering Department, starting with one lab principal investigator (PI: Macklin), one scientific staff (Heiland), and one Ph.D. student. While the PI had previously taught dedicated classes on mathematical and computational biology and had involved undergraduate students in research, these efforts were performed separately. The move to a new program presented a rare opportunity to build and iteratively refine a new research program that involved undergraduate students from the very start. In the Fall semester, the lab began integrating undergraduate researchers into its growing research program, with several guiding principles:

- Students should be involved with the main research program. (Students should be directly involved in ongoing publication-driven research, rather than projects created solely for didactic purposes.)
- Student involvement should accelerate these existing projects or allow expanded exploration of the existing scientific aims.
- More experienced students should help mentor less experienced students to foster a sustainable team.
- Research results should feed back into education and outreach.
- Graduate students should gain team management experience while helping to mentor undergraduate students.
- The students’ class work and personal/home responsibilities must take priority over their research involvement.
- Students should be encouraged to seek other opportunities in the summer to broaden their skills and drive professional networking.

### 2.1 Evolving undergraduate research model

Motivated by rapid prototyping methodologies, we iteratively developed, tested, and refined the mentorship structure (with guidance from computational apprenticeship theory) while integrating undergraduate students into ongoing (primarily grant-funded) research in mathematical oncology. At the end of each semester, we evaluated the current mentorship structure against the guiding principles, with particular attention to:

1. progress towards research milestones
2. progress towards peer-reviewed scientific posters
3. development of scientific and communication skills
4. individual understanding of the projects and their contributions
5. unanticipated creativity and innovation
6. emergence of undergraduate student leadership
7. evolving team leadership skills by involved graduate students
8. undergraduate student retention

The progress was assessed by a combination of graduate student, one research staff (Heiland), and PI (Macklin) observations, student interviews (lead by Madamanchi [48]), and end-of-semester lab discussions.

### 2.2 Version 1 (Fall 2017)

The first version of the mentorship structure included the PI (Macklin), one scientific research staff (Heiland), one Ph.D. student, and five undergraduate (freshman) students with a variety of backgrounds in engineering and neuroscience. See Table 1 in APPENDIX A.

**Table 1.** Evolving MathCancer lab and mentoring structure.

|  | Version 1 | Version 2 | Version 3 | Version 4 |
| --- | --- | --- | --- | --- |
| Scientific Staff | 1 | 1 | 1 | 1 |
| Ph.D. Students | 1 | 3 | 3 | 5 |
| Undergraduate Trainees | 5 | 6 | 10 | 10 |
| Fields | engineering neurobiology | engineering | engineering CS, informatics | engineering CS, informatics |
| Number of Teams / Projects | 1 | 3 | 5 | 3 |
| Co-mentors? |  | Yes | Yes | Yes |
| Weekly meeting? |  | Yes | Yes | Yes |
| State of the Lab? |  | Yes | Yes | Yes |
| Mixed update and skills talks |  |  |  | Yes |

This mentoring structure focused on training the undergraduate and graduate students to use the lab’s main computational framework (Physi-Cell [40]). Each week, the group met for a 1-2 hour live coding session that introduced the codebase and illustrated modeling techniques for sample tumor growth problems. The Ph.D. student and research staff attended the sessions and helped the undergraduate researchers to troubleshoot their code, similarly to the role of teaching assistants in lab sections for programming-heavy STEM courses.

#### Assessment

At the end of the semester, the PI, scientific staff, and graduate student met to discuss the successes and failures of the semester, based on their personal observations.

#### Research impact

These coding sessions exposed areas for improvement in PhysiCell, particularly ways that model setup could be automated and made more user-friendly. These core method improvements were beneficial to all scientific projects in the lab.

#### Other metrics

One of the five students returned to continue research in the following semester.

#### What worked

The students progressed from little-to-no programing expertise, to being able to independently compile and run C++-based PhysiCell simulations on their own laptops. They learned how to make minor code modifications to existing models to change the model hypotheses, often driven by a basic understanding of ordinary differential equations. They also learned to create and present scientific posters.

#### Areas for improvement

We found that the live coding sessions did not make the fullest use of the students’ individual capabilities. The students needed more hands-on time to learn and contribute individually. The computational apprenticeship principle of ‘fading’, in which mentors gradually give less support, needs more time to implement.

### 2.3 Version 2 (Spring 2018)

In response to the Version 1 observations, we changed to a small team structure. Each team consisted of 1-3 undergraduate students, the PI, and potentially a co-mentor (Ph.D. student or scientific staff). Each team met early in the week for approximately 1 hour to mentor, set goals, and work. The undergraduate students worked on their own towards the weekly goals for 1-2 hours between these weekly mentored meetings. The updated lab structure is in Table 1 in APPENDIX A. Note that all the undergraduate students were freshmen. The semester’s projects included:

**Project 1:** Develop Jupyter notebook user interfaces for PhysiCell models (PI, research staff, 3 undergraduates).

**Project 2:** Develop a model of extracellular matrix (ECM) remodeling by migrating tumor cells (PI, Ph.D. student, 2 undergrads).

**Project 3:** Develop a model of colon cancer metastases (PI, 1 undergraduate).

#### Assessment

At the end of the semester, the entire lab met to discuss the semester’s progress and assess our current research organization. Afterwards, the PI, scientific staff, and Ph.D. student met to discuss the final lab meeting’s observations, together with their own personal observations. Madamanchi began collaborating in Summer 2018 to observe the lab structure and contribute computational apprenticeship expertise to help refine our mentoring structure [48].

#### Research impact

The students in Project 1 were successful in prototyping a technique to create a Jupyter-based graphical user interface (GUIs) for a PhysiCell-based simulation model of cancer nanotherapy [49]. They presented their work at an Indiana University poster session and at a major NSF site visit. The students in Project 2 were able to prototype key elements of the ECM model and present their results at a poster session. The student in Project 3 was not able to make progress, primarily due to other extracurricular priorities for the student.

#### Other metrics

Five of the six students returned to continue research in the following semester. The remaining student was not asked to return due to insufficient research progress.

#### What worked

The students were able to make individual contributions to the projects. We observed growth in their C++ and Python skills, and independent creativity (particularly in Project 1).

#### Areas for improvement

While the students were able to make individual contributions to their projects, they expressed that they felt isolated from the lab and unaware of progress by other teams. They sought increased interactions between the teams. This is consistent with computational apprenticeship, which emphasizes the need for social learning and peer-to-peer information transfer. The PI observed that his weekly meetings with each team were not scalable or sustainable.

### 2.4 Version 3 (Fall 2018–Spring 2019)

To address our Version 2 observations, we refined the mentoring structure to ensure that each team had a non-PI co-mentor who would take responsibility for the team. The updated lab structure is in Table 1 in APPENDIX A. We also altered the weekly mentoring schedule:

- The PI met with all co-mentors in weekly one-on-one mentoring to discuss their team progress and plan their team’s next steps.
- The non-PI co-mentors met with their teams early each week (approximately 1 hour) to set goals and work. The PI attended these meetings by request of the co-mentors.
- The undergraduate researchers worked on their own towards the weekly goals mid-week and/or on the weekend.
- We held an “all hands” lab meeting each Friday for 1-2 hours:

> **Team presentation:** One of the teams prepares and presents a 10-20 minute presentation on their progress and open problems, followed by group discussion. This encouraged “cross-pollination” between teams and collective brainstorming, while developing undergraduate student presentation skills and encouraging individual student understanding of the work.
>
> **Unstructured mentoring time:** For the remainder of the group meeting, we broke into teams, while the PI met with each team for extra mentoring and troubleshooting.
- We added a PI’s “state of the lab” talk to the end of each semester to summarize and contextualize progress and kick-start group discussion to assess our lab processes.

In these semesters, the projects included:

**Project 1:** Continue development of Jupyter notebook user interfaces for computational models (research staff, 3 undergraduates)

**Project 2:** Continue development of the ECM model (Ph.D. student 1, 3 undergraduates),

**Project 3:** Develop an improved nanoparticle model (Ph.D. student 2, 1 undergraduate)

**Project 4:** Continue developing of PI’s prototype of a cancer hypoxia model (Ph.D. student 3, 2 undergraduates)

**Project 5:** Extend software for Microsoft Windows compatibility (research staff, 1 undergraduate)

#### Assessment

At each semester’s final all-hands meeting, the entire lab (PI, scientific staff, Ph.D. students, undergraduate researchers) discussed the semester’s progress and name strengths and weaknesses of our current research organization. Afterwards, the PI, scientific staff, and Ph.D. students met to discuss the final lab meeting’s observations, together with their own personal observations. Madamanchi performed student interviews (results were published in [48]) and consulted regularly with the PI on his observations.

#### Research impact

The students in Project 1 were successful in generalizing their previous prototype to develop xml2jupyter, which was featured in a student co-authored scientific abstract and peer-reviewed scientific paper [50]. Students in Projects 2, 3 and 4 continued to refine their computational models and presented their owrk at a poster session. Projects that had new undergraduate students and graduate mentors had more limited progress as those students needed more extensive PI training. The student in Project 5 made substantial progress in their software development project [51]. Two additional undergraduate students attended lab meetings irregularly and contributed to discussions but did not attend regularly enough to join projects.

#### Other metrics

One undergraduate student served as an undergraduate team lead in Project 2 and began training a successor so he could pursue interests in his core concentration (cyberphysical systems and computer engineering). Three undergraduate students contributed to a peer-reviewed journal article [50]; two of these ramped down their involvement and “graduated” from the lab after reaching this milestone, allowing them to pursue interests in their concentration of study. One student graduated from Indiana University and was employed in industry. Five of the remaining students returned to continue research in the following academic year. (As noted above, three left the group to research closer to their engineering concentration after successful knowledge transfer, and one graduated.)

#### What worked

The students made substantive individual contributions, including a peer-reviewed publication [50]. Student creativity lead to unanticipated advances (*ad hoc* small team crowdsourcing), and we observed frequent undergraduate-undergraduate mentoring. Notably, this updated mentoring structure accommodated a near doubling of undergraduate involvement (an increase from six to ten students).

#### Areas for improvement

Overall, we have found that this mentoring structure has been successful, but we identified areas for improvement. The large number of projects left the lab feeling fragmented and difficult to manage. Some of the teams were unbalanced: Projects 1, 2, and 5 benefited from a senior Ph.D. student (with prior mentoring experience in industry) or scientific staff. Projects 3-4 were co-mentored by younger Ph.D. students with less leadership experience and domain knowledge, leading to reduced progress. Student surveys also found that students in Projects 3-4 gained the impression that Projects 1-2 and 5 were higher lab priorities. Thus, better communication of priorities and the impact of each project were needed in the younger teams.

### 2.5 Version 4 (Fall 2019–present)

The Version 4 lab structure is in Table 1 in APPENDIX A. We modified the Version 3 mentoring structure in several ways:

- Organize teams around themes, rather than specific technical projects:

**Team 1:** <u>PhysiCell training and community</u>: In support of a new NCI administrative supplement, this team worked to develop interactive training materials red: a series of 10-15 minute training modules including PowerPoint slides, YouTube recordings with captions, and nanoHUB-hosted microapps to illustrate core code concepts. They also worked on developing a website to focus the growing international PhysiCell modeling community. (2 Ph.D. students, 4 undergraduate students)

**Team 2:** <u>ECM model development</u>: This team performed final computational investigations of the ECM model and worked on a scientific manuscript. The group also worked on drafting a scientific manuscript for the *ad hoc* crowdsourcing technique. (1 Ph.D. student, 2 undergraduate students)

**Team 3:** <u>PhysiCell tools</u>: This team continued xml2jupyter [50] refinements, but was also encouraged to creatively explore standalone tools that could increase the usability and utility of PhysiCell models. (1 staff, 2 Ph.D. students, 4 undergraduate students)

- Devote some lab presentations to new team management or technical skills, rather than project progress. Examples included:
  - Project management with Trello
  - Scrums, sprints, and kanbans (software team skills)
  - PhysiCell simulation data structures [46]

- The PI gives frequent updates from research travel and reinforces the key role of each team’s work in the lab’s long-term strategy.

#### Assessment

This iteration is still in progress. An end-of-semester “state of the lab” talk and discussion is planned for December 2019, as well as discussion among the senior staff (Ph.D. students, scientific staff, and PI) and Madamanchi. Macklin and Madamanchi are planning assessments for use in the Spring 2020 semester.

#### Research impact

Team 1 has prototyped educational microapps and tested presentations. They began brainstorming new outreach methods while designing the PhysiCell.org website. Team 2 continues to make good progress on their ECM model and is performing final analyses for their manuscript. Team 3 has released a “Python loader” tool [52, 53] to load simulation data into Python, and add significant visualization and usability refinements.

#### Other metrics

This work is ongoing, but we see evidence of strong student leadership. The students in Team 1 (mostly sophomores) developed their own recording methodologies and are leading the development of educational microapps. They have also proposed leading a student-run minisymposium at the 2020 Annual Meeting of the Biomedical Engineering Society (BMES), showing a sense of intellectual co-ownership. Team 2 is working independently: the PI coordinates work with the Ph.D. student lead, who has also encouraged leadership by the undergraduate students. Team 3 has shown substantial technical know-how and creativity in adapting open source tools to the PhysiCell software ecosystem. They are actively leading the development of new features.

#### What worked

While this lab version is ongoing, preliminary observations find that grouping more students together in teams wrapped around themes has allowed greater peer-to-peer mentoring, individual creativity, and initiative. The students frequently suggest solutions and meet in smaller pairs to work new angles. Teams 1 and 3 have begun breaking their topics down into separate sub-projects to work in parallel.

Pairing two younger Ph.D. students in Team 1 was helpful in addressing the prior weakness of unbalanced teams (particularly teams where the undergraduate and graduate students were less experienced). Moreover, mixing new and returning students in Teams 1 and 3 helped to balance expertise and encourage within-team peer mentoring.

#### Areas for improvement

Overall, we have found that this mentoring structure has addressed most of the issues identified in Version 3. We will continue to evaluate and refine.

### 2.6 Contextualization as Computational Apprenticeship

Undergraduate research is broadly considered a valuable component of undergraduate education in STEM disciplines. Participation in undergraduate research is associated with greater STEM retention and educational achievement [54–56]. However, the traditional models for undergraduate research have developed in the natural sciences, and there is a need for critical reflection on how to scale and adapt undergraduate research opportunities within computational and interdisciplinary fields.

The MathCancer lab’s evolving approach to undergraduate mentoring has developed a tiered mentoring structure that aligns with education research showing that students benefit from having both faculty and graduate or staff mentors [57, 58]. The apprenticeship relationship between undergraduate researchers and their mentors provides not only domain knowledge, but also higher-order skills including heuristic strategies, learning strategies, and metacognitive skills that are associated with ‘thinking like a scientist’ [59]. Additionally, undergraduate researchers are socialized into the disciplinary community and can have gains in their disciplinary identity that are associated with long-term persistence within the field [60]. A tiered mentoring structure allows each undergraduate researcher to receive coaching and guidance from multiple mentors, as well as provides opportunities for students to learn by observing and modeling the disciplinary processes of their mentors and their undergraduate peers. Similarly, structured interactions with multiple lab members supports the integration of the undergraduate researchers in the lab community. At primarily undergraduate institutions in which staff or graduate secondary mentors are not possible, more advanced undergraduates can serve many of these important functions.

The MathCancer group recently partnered with computational education researchers to study the experience of their undergraduate researchers [48]. Our qualitative investigation found that the undergraduate researchers enjoyed and valued their time in the MathCancer group. Specifically, the students reported gains in their intellectual and personal development through their lab experience. Students reported increased knowledge of “real-world” engineering and modeling norms and practices, and they reported gains in their ability to “think like an engineer”, suggesting growth in both metacognitive skills and disciplinary identity. The students’ self-reported development in these domains is similar to published findings from studies of undergraduate researchers in the natural sciences [60–62].

Our study also characterized the executive management and strategic knowledge component of the undergraduate researchers’ metacognition. Students displayed varied levels of these metacognitive skills, which reflected the differences in age and length of research experience. The students all reported an executive management approach of “guess and check” for implementing their research plan. More mature students were able to identify this approach as part of a larger iterative process of planning and evaluating. In contrast, students with less experience in the lab indicated a high degree of reliance upon their staff or graduate co-mentor to help identify the next step in the plan. The tiered mentorship structure of the MathCancer research group allows for greater scaffolding and support for novice researchers and builds in “fading” of that support and greater independence for more advanced undergraduates. The undergraduate researchers all indicated satisfaction with the growth in their strategic knowledge, but had difficulties in articulating the heuristic, control, and learning strategies that they use in the research process. Interviews with faculty, staff, and graduate mentors as well as examination of the group’s research documentation suggests that the undergraduate researchers did, in fact, gain experience with new heuristics for problem-solving but had difficulty recalling or articulating them in a decontextualized semi-structured interview.

To further support the metacognitive development of its undergraduate researchers, the MathCancer group intends to provide mentorship training to lab members and embed scaffolded reflection to its research process. Mentorship training will consist of a short seminar on computational apprenticeship model. This seminar is intended to remind both the mentors and the undergraduate researchers that learning consists not only of technical domain knowledge, but also of metacognitive knowledge. The seminar will also cover the modes and sequencing of mentorship to help mentors understand different ways of organizing research tasks, and prompt students to be more intentional about their learning process (see Table 2 in APPENDIX B, constructed from [30, 59]). The mentors in the MathCancer group already demonstrate many of the mentoring modalities of computational apprenticeship, but research indicates that foregrounding these approaches through mentorship training is beneficial for undergraduate researchers [63, 64].

**Table 2.** Computational Apprenticeship: Mode and Sequencing of Mentoring.

| **Mode of Mentoring** |  |
| --- | --- |
| Modeling: | Explicit demonstration of a task, including verbalizing the associated heuristics (strategies) |
| Coaching: | Observing students as they perform tasks and offering feedback |
| Scaffolding: | Making tasks accessible to students by calibrating difficulty levels |
| Articulation: | Asking students to verbalize their process as they complete tasks |
| Reflection: | Prompting students to compare multiple approaches to problem solving |
| Exploration: | Fading or slowly withdrawing as students gain the ability to perform complex tasks |
| **Sequencing of Mentoring** |  |
| Increasing complexity: | organizing coding tasks from simple to more complex |
| Increasing diversity: | allowing students to develop skills within one language/project before transferring those approaches to a new context |
| Global to local skills: | Sharing the overall conceptual approach using psuedocode before implementing specific sub-tasks |

Similarly, the MathCancer group plans to embed bimonthly scaffolded reflection prompts into their existing research documentation process. Specifically, they have adapted Howitt et al.’s Learner Logbook intervention for computational research as a way helping undergraduates absorb bigger picture learning during their research [65]. See Table 3 in APPENDIX B.

**Table 3.** Scaffolded Reflection Prompts.

|  |  |
| --- | --- |
| Students will select one question to briefly answer every two months: |  |
| • | How has your group navigated any challenges you have encountered? |
| • | What might you have done differently if you had known two months ago what you know now? |
| • | Has your research question changed? If so, why, and what has it changed to? |
| • | How have you chosen the approach or methods that you are using for your project? |
| • | What are the connections between your research activities and your other studies? |
| • | Can you see ways in which you could apply what you have learned to other activities? |

## 3 Ongoing and future work: extending computational apprenticeship to classroom instruction and scholarly communication

For many computational research labs, the educational infrastructure of the undergraduate research experience is often considered to be distinct from the software infrastructure developed by the lab. However, there are key parallels: as a platform that facilitates computational modeling, PhysiCell provides ‘software-realized scaffolding’ that enables scientists to more easily engage with disciplinary biology questions within the computational realm [66, 67]. Students with moderate programming experience can extend sample projects to implement complex models that integrate cutting-edge stochastic processes, 3-D diffusion, discrete processes without building these capabilities “from scratch,” allowing them to focus more deeply on (1) “translating” biological behaviors into mathematical submodels and agent rules, (2) rapidly testing and visualizing the emergent nonlinear behaviors of these complex systems, (3) analyzing their results, and (4) communicating what they have learned. Interdisciplinary undergraduate instruction, and scientific communication are two arenas in which the software infrastructure of PhysiCell can support a computational apprenticeship analogous to the undergraduate research program of the MathCancer Group.

A major goal of interdisciplinary education at the intersection of life sciences and computational sciences is acculturating students to the modes of thinking within each discipline. Authentic learning experiences that provide students with realistic interdisciplinary problems are crucial for teaching students to “think like a biologist” and “think like an engineer/mathematician”.

However, presenting students with computational or systems concepts applied to life sciences problems can pose challenges to students by simultaneously introducing new disciplinary knowledge and new technical content. The use of educational “microapps” or interactives can be a powerful way of creating authentic learning experiences, while still providing the scaffolding and sequencing called for within the computational apprenticeship framework. Microapps can embed widgets and sliders that allow students to explore the problems globally or conceptually before implementing models on their own. Moreover, interactive exploration of complex computational models could help students build intuition for deep mathematical concepts far earlier than they are introduced in most classroom-based curricula. We found that freshmen researchers gained intuition in stochastic processes (e.g., Poisson birth-death processes and biased random walks) and partial differential equations (diffusion processes, Dirichlet and von Neuman boundary conditions, diffusion length scales) from using and developing these computational models. Future work should focus on how these apps can be refined and augmented with video recording and other scaffolding to improve this learning.

The MathCancer group is adapting xml2jupyter [50] (see Section 2.4) to build technological infrastructure to support these pedagogical approaches. Small, efficient agent-based models [41] can be purpose-built to illustrate biological concepts, automatically fitted with Jupyter-based GUIs, and rapidly deployed as cloud-hosted educational microapps. These apps present didactic information, parameter tabs (default values guide novice users), a runnable simulation model, and tabs to visualize the cell behavior and chemical substrates. (See Figure 2 for an example that explores biased random cell migration.) Macklin has tested microapps in an undergraduate systems biology course, and the MathCancer lab is using the approach to build a series of microapp-enhanced training modules for PhysiCell. (See Section 2.5.) Moreover, Macklin uses the xml2jupyter workflow to enhance computational apprenticeship in the classroom: advanced multicellular systems biology students at Indiana University develop their own PhysiCell [40] models as a final project, convert them to cloud-hosted models with xml2jupyter [50], and demonstrate their interactives in their final presentations. Future PhysiCell educational microapps could be developed, in careful collaboration with instructional experts, to supplement open educational materials in educational communities such as QUBES [68]. Moreover, such educational communities could act as “educational marketplaces,” connecting tool builders (e.g., engineering faculty) with educators including biologists and mathematicians who use interactives in their curriculum. Such partnerships between modelers and educators are critical in ensuring that shared research models are useful to student learning with sufficient scaffolding. This is an increasing area of focus for the MathCancer lab, and early feedback informs us that a scaffolding with the background mathematics, expected model behavior, and suggested exploration improves student experiences with these models. We are currently exploring the use of online videos on specific apps as further scaffolding.

**Figure 2.**
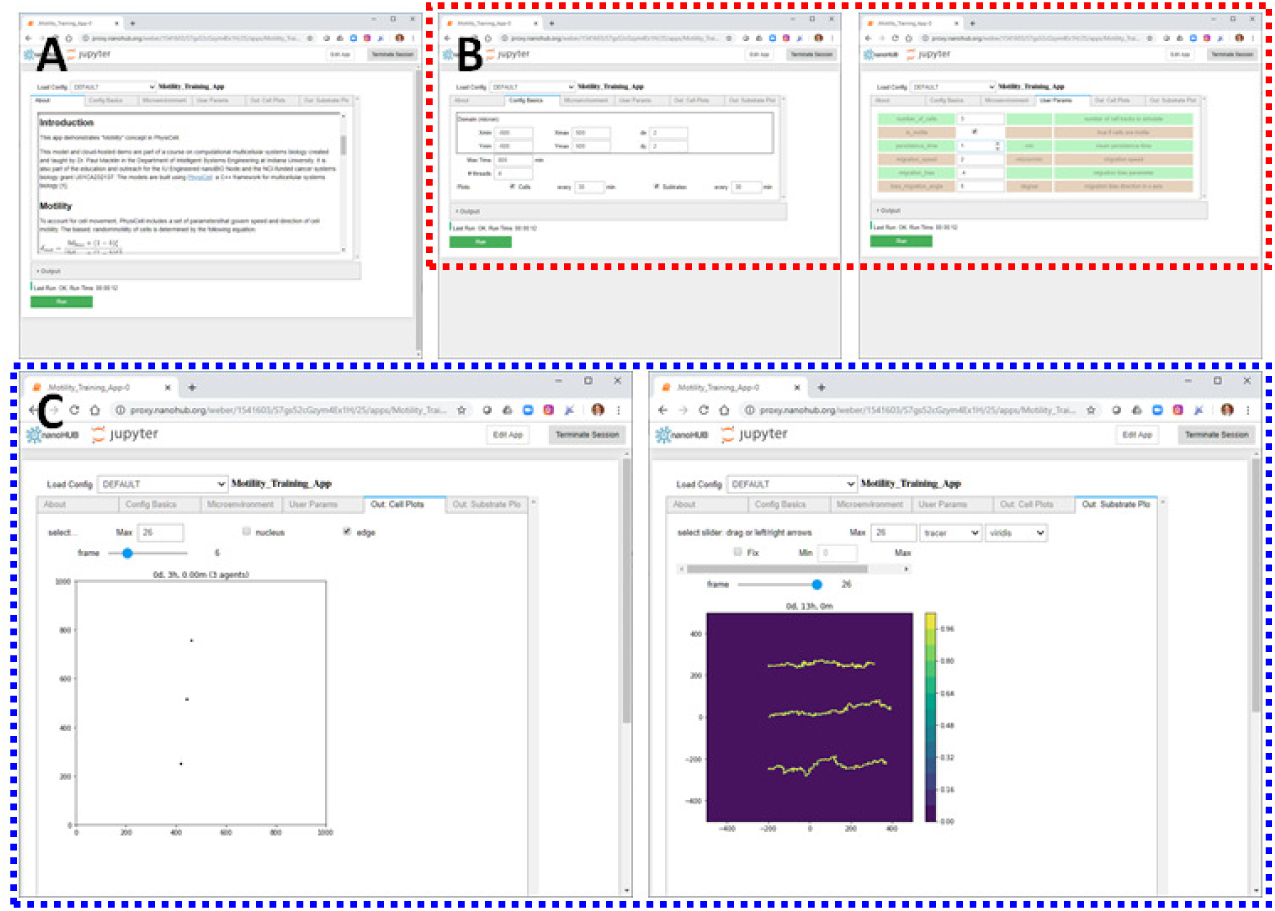
Educational microapps. The first PhysiCell/xml2jupyter educational microapp, [69], was rapidly developed and deployed over 2 days in response to student learning needs. As part of Team 1’s work in the Version 4 lab (2.5), this has been refined to build a new microapp ([70]) to illustrate biased random cell migration in PhysiCell in train users to set key phenotypic parameters. Note that the app has didactic material (A), user-set parameters (B), a runnable simulation, and built-in visualization (C).

Similar approaches may be valuable for interdisciplinary communication among practicing scientists. Current practices for communicating computational biology are limited to traditional paper formats that include the foundational equations or at best a link to the model code. This approach puts a high burden for replication and exploration on time-limited readers. Worse, for biologists with limited computational training, this approach prevents any engagement with computational biology literature. The MathCancer Lab used xml2jupyter [50] to create pc4cancerimmune [43], its first “publication companion app” as part of [38]. This allows scientific readers to interactively explore and understand the key cancer immunology simulation model at the heart of the publication’s method, as well as to better disseminate model to the broader scientific community. Moreover, cloud-hosted versions of published research-grade models can readily be used in classroom instruction, thus allowing educators to rapidly incorporate cutting-edge research in their curriculum. The MathCancer lab has tested using publication companion apps to illustrate intelligent systems modeling to sophomore engineering students at Indiana University, and it is currently seeking new educational communities to further test the concept.

The current work and future directions of the MathCancer lab highlight the value and mutual benefit of engaging with education scholarship and education researchers. We have observed several instances of what may be termed ‘convergent evolution’ between our practice and education research findings. Our hierarchical mentoring model includes the mentoring triads that education researchers have identified as a best practice. Similarly, our approach to ‘rapid prototyping’ our educational infrastructure mirrors the design-based research modality of education researchers. Consultation with education researchers can help educators to arrive at theoretically and empirically supported practices more quickly. Similarly, practitioners in rapidly evolving interdisciplinary education spaces can offer new perspectives that stimulate new education scholarship. Direct collaboration with the educational community will be essential if research-driven apps are to reach their pedagogical potential.

## ACKNOWLEDGMENTS

Please see the title page (after blinded review).

## APPENDICES

### APPENDIX A MathCancer Lab Mentoring Structure

We summarize and compare the Version 1 (Section 2.2), Version 2 (Section 2.3), Version 3 (Section 2.4), and Version 4 (Section 2.5) lab structure versions in Table 1.

### APPENDIX B Computational Apprenticeship mentorship & reflection

Modes and sequencing of mentoring in Computational Apprenticeship are given in Table 2. Scaffolded reflection prompts are given in Table 3

## BIOGRAPHICAL SKETCHES

Aasakiran Madamanchi completed his doctorate in Cancer Biology at Vanderbilt University. He is now a Postdoctoral Associate in the Future Work and Learning Impact Area in the Polytechnic Institute at Purdue University. His primary research interest is in democratizing computational modeling and data science approaches. He is particularly interested in promoting and supporting computational modeling in the life sciences.

Madison Thomas is an undergraduate at Purdue University graduating with her Bachelors in Computer and Information Technology in May 2020. She enjoys working with students and researching topics related to computing.

Alejandra Magana completed her graduate education in engineering education at a midwestern research university where she is now a professor of computer and information technology. Her research program seeks to understand how computational model-based reasoning can effectively support scientific inquiry learning, problem solving, and innovation processes in the context of authentic modeling and simulation practices.

Randy Heiland received degrees in computer science (Utah) and mathematics (Arizona State) and has pursued opportunities to effectively combine them ever since, including numerous education and outreach opportunities in K-16. A simple recipe he has followed is to use age-appropriate mathematics with interactive computer simulations to get young (and not so young) people excited about science and math.

Paul Macklin studied mathematics at the University of Nebraska-Lin-coln, the University of Minnesota, and the University of California-Irvine. He has developed in open source software for mathematical oncology with academic positions in mathematics, health informatics, research medicine, and engineering. He is an Associate Professor and Director of Undergraduate Studies in Intelligent Systems Engineering at Indiana University. He seeks to involve undergraduate students in cutting-edge research and bring that research to the classroom.

## REFERENCES

[1] A A Seyhan. Lost in translation: the valley of death across preclinical and clinical divide – identification of problems and overcoming obstacles. Transl. Med. Commun., 4(18):1–19. doi: 10.1186/s41231-019-0050-7. URL https://doi.org/10.1186/s41231-019-0050-7.

[2] Jack W Scannell, Alex Blanckley, Helen Boldon, and Brian Warrington. Diagnosing the decline in pharmaceutical R&D efficiency. Nat. Rev. Drug Disc., 11(3):191, 2012. doi: 10.1038/nrd3681. URL https://dx.doi.org/10.1038/nrd3681.

[3] Jiaquan Xu, Sherry L Murphy, Kenneth D Kochanek, Brigham Bastian, and Elizabeth Arias. Deaths: final data for 2016. National vital statistics reports, 67(5), 2018. URL https://stacks.cdc.gov/view/cdc/57989.

[4] Phillip A Sharp and Robert Langer. Promoting convergence in biomedical science. Science, 333(6042):527–527, 2011. doi: 10.1126/science.1205008. URL https://dx.doi.org/10.1126/science.1205008.

[5] National Research Council. Convergence: facilitating transdisciplinary integration of life sciences, physical sciences, engineering, and beyond. National Academies Press, 2014. ISBN 9780309301640. doi: 10.17226/18722. URL https://dx.doi.org/10.17226/18722.

[6] Jonathan Ozik, Nicholson Collier, Justin Wozniak, Charles Macal, Chase Cockrell, Samuel H Friedman, Ahmadreza Ghaffarizadeh, Randy Heiland, Gary An, and Paul Macklin. High-throughput cancer hypothesis testing with an integrated PhysiCell-EMEWS workflow. BMC Bioinformatics, 19(Suppl 18):483, 2018. doi: 10.1186/s12859-018-2510-x. URL http://dx.doi.org/10.1186/s12859-018-2510-x.

[7] Paul Macklin. Key challenges facing data-driven multicellular systems biology. GigaScience, 8(10), 2019. doi: 10.1093/gigascience/giz127. URL http://dx.doi.org/10.1093/gigascience/giz127.

[8] Brenden K. Petersen, Jiachen Yang, Will S. Grathwohl, Chase Cockrell, Claudio Santiago, Gary An, and Daniel M Faissol. Deep reinforcement learning and simulation as a path toward precision medicine. J. Comput. Biol., 26(6):597–604, 2019. doi: 10.1089/cmb.2018.0168.

[9] Michael Pargett, Ann E Rundell, Gregery T Buzzard, and David M Umulis. Model-based analysis for qualitative data: an application in drosophila germline stem cell regulation. PLoS Comput. Biol., 10(3):e1003498, 2014. doi: 10.1371/journal.pcbi.1003498. URL https://dx.doi.org/10.1371/journal.pcbi.1003498.

[10] Aasakiran Madamanchi, Monica E Cardella, James A Glazier, and David M Umulis. Factors mediating learning and application of computational modeling by life scientists. In 2018 IEEE Frontiers in Education Conference (FIE), pages 1–5. IEEE, 2018. doi: 10.1109/fie.2018.8659328. URL https://dx.doi.org/10.1109/fie.2018.8659328.

[11] Aasakiran Madamanchi, Shenika V Poindexter, Monica E Cardella, James A Glazier, and David M Umulis. Qualitative findings from study of interdisciplinary education in computational modeling for life sciences student researchers from emerging research institutions. In 2018 IEEE Frontiers in Education Conference (FIE), pages 1–5. IEEE, 2018. doi: 10.1109/FIE.2018.8659183. URL https://dx.doi.org/10.1109/FIE.2018.8659183.

[12] William Bialek and David Botstein. Introductory science and mathematics education for 21st-century biologists. Science, 303 (5659):788–790, 2004. doi: 10.1126/science.1095480. URL https://dx.doi.org/10.1126/science.1095480.

[13] Pat Marsteller, Lisette de Pillis, Ann Findley, Karl Joplin, John Pelesko, Karen Nelson, Katerina Thompson, David Usher, and Joseph Watkins. Toward integration: from quantitative biology to mathbio-biomath? CBE—Life Sciences Education, 9(3):165–171, 2010. doi: 10.1187/cbe.10-03-0053. URL https://dx.doi.org/10.1187/cbe.10-03-0053.

[14] Pat Marsteller. Beyond bio2010: integrating biology and mathematics: collaborations, challenges, and opportunities. CBE life sciences education, 9(3):141, 2010. doi: 10.1187/cbe.10-06-0084. URL https://dx.doi.org/10.1187/cbe.10-06-0084.

[15] Jason Feser, Helen Vasaly, and Jose Herrera. On the edge of mathematics and biology integration: improving quantitative skills in undergraduate biology education. CBE—Life Sciences Education, 12(2):124–128, 2013. doi: 10.1187/cbe.13-03-0057. URL https://doi.org/10.1187/cbe.13-03-0057.

[16] Eroma Abeysinghe, Michal Brylinski, Marcus Christie, Suresh Marru, and Marlon Pierce. Lsu computational system biology gateway for education. In Proceedings of the Practice and Experience in Advanced Research Computing on Rise of the Machines (learning), pages 1–4. 2019.

[17] Paul A Craig. Developing and applying computational resources for biochemistry education. Biochemistry and Molecular Biology Education, 2020.

[18] Anna Ritz. Programming the central dogma: An integrated unit on computer science and molecular biology concepts. In Proceedings of the 49th ACM Technical Symposium on Computer Science Education, pages 239–244, 2018.

[19] Tanya Berger-Wolf, Boris Igic, Cynthia Taylor, Robert Sloan, and Rachel Poretsky. A biology-themed introductory cs course at a large, diverse public university. In Proceedings of the 49th ACM Technical Symposium on Computer Science Education, pages 233–238, 2018.

[20] Arthur T Johnson. Biology for engineers. CRC Press, 2011. ISBN 9781138067899.

[21] William H O Guilford. A problem-based approach to teaching cell and molecular biology to engineers. Proceedings of the 2001 American Society for Engineering Education Annual Conference & Exposition, 6:1, 2001. URL https://peer.asee.org/9676.

[22] C Ramey, Judy Schoonmaker, and S Ryan. Studio biology for engineers: Lessons learned. In 2017 ASEE Annual Conference & Exposition, Columbus, Ohio, 2017. URL https://peer.asee.org/28874.

[23] Alejandra J Magana, Manaz Taleyarkhan, Daniela Rivera Alvarado, Michael Kane, John Springer, and Kari Clase. A survey of scholarly literature describing the field of bioinformatics education and bioinformatics educational research. CBE—Life Sciences Education, 13(4):607–623, 2014. doi: 10.1187/cbe.13-10-0193. URL https://dx.doi.org/10.1187/cbe.13-10-0193.

[24] Hayden Fennell, Joseph A. Lyon, Aasakiran Madamanchi, and Alejandra J Magana. Computational apprenticeship: Cognitive apprenticeship for the digital era. SocArXiv, 2019. doi: 10.31235/osf.io/jy328. URL https://dx.doi.org/10.31235/osf.io/jy328.

[25] Matthew Butler, Michael Morgan, et al. Learning challenges faced by novice programming students studying high level and low feedback concepts. In Proceedings ascilite Singapore, number 99–107, 2007. URL http://www.ascilite.org/conferences/singapore07/procs/butler.pdf.

[26] AM Heroux, G Allen, et al. Computational science and engineering software sustainability and productivity (csessp) challenges workshop report. Arlington, VA: Networking and Information Technology Research and Development (NITRD) Program, 2016. URL https://www.nitrd.gov/pubs/CSESSPWorkshopReport.pdf.

[27] Camilo M Viera, Alejandra J Magana, Anindya M Roy, Michael L Falk, Michael J Reese Jr, et al. Exploring undergraduate students’computational literacy in the context of problem solving. The ASEE Computers in Education (CoED) Journal, 7(1):100, 2016. doi: 10.18260/p.24081. URL https://dx.doi.org/10.18260/p.24081.

[28] Huma Shoaib, Monica Cardella, Aasakiran Madamanchi, and David Umulis. An investigation of undergraduates’ computational thinking in a sophomore-level biomedical engineering course. Proceedings–Frontiers in Education Conference, FIE, October 2019, Covington, KY, USA, pages 1–4, 2019. doi: 10.1109/FIE43999.2019.9028503.

[29] Allan Collins, John Seely Brown, and Ann Holum. Cognitive apprenticeship: Making thinking visible. American educator, 15(3): 6–11, 1991.

[30] Allan Collins, John Seely Brown, and Susan E Newman. Cognitive apprenticeship: Teaching the craft of reading, writing and mathematics. Thinking: The Journal of Philosophy for Children, 8(1): 2–10, 1988.

[31] Paul Macklin. When Seeing Isn’t Believing: How Math Can Guide Our Interpretation of Measurements and Experiments. Cell Sys., 5 (2):92–4, 2017. doi: 10.1016/j.cels.2017.08.005. URL http://dx.doi.org/10.1016/j.cels.2017.08.005.

[32] Paul Macklin, Hermann B Frieboes, Jessica L Sparks, Ahmadreza Ghaffarizadeh, Samuel H Friedman, Edwin F Juarez, Edmond Jockheere, and Shannon M Mumenthaler. Progress towards computational 3-d multicellular systems biology. In K A Rejniak, editor, Systems Biology of Tumor Microenvironment, chapter 12, pages 225–46. Springer, 2016. doi: 10.1007/978-3-319-42023-312. URL https://dx.doi.org/10.1007/978-3-319-42023-3_12. (invited author: P. Macklin).

[33] Ariel L. Rivas, Gabriel Leitner, Mark D. Jankowski, Almira L. Hoogesteijn, Michelle J. Iandiorio, Stylianos Chatzipanagiotou, Anastasios Ioannidis, Shlomo E. Blum, Renata Piccinini, Athos Antoniades, Jane C. Fazio, Yiorgos Apidianakis, Jeanne M. Fair, and Marc H. V. Van Regenmortel. Nature and consequences of biological reductionism for the immunological study of infectious diseases. Frontiers in Immunology, 8:612, 2017. doi: 10.3389/fimmu.2017.00612. URL https://www.frontiersin.org/article/10.3389/fimmu.2017.00612.

[34] Fulvio Mazzocchi. Complexity and the reductionism–holism debate in systems biology. Wiley Interdisciplinary Reviews: Systems Biology and Medicine, 4(5):413–427, 2012. doi: 10.1002/wsbm.1181. URL https://onlinelibrary.wiley.com/doi/abs/10.1002/wsbm.1181.

[35] Gunver Kienle and Helmut Kiene. From reductionism to holism: Systems-oriented approaches in cancer research. Global Advances in Health and Medicine, 1(5):68–77, 2012. doi: 10.7453/gahmj.2012.1.5.015. URL https://doi.org/10.7453/gahmj.2012.1.5.015. PMID: 27257534.

[36] Jeffrey West and Paul K. Newton. Cellular interactions constrain tumor growth. Proceedings of the National Academy of Sciences, 116(6):1918–1923, 2019. doi: 10.1073/pnas.1804150116. URL https://www.pnas.org/content/116/6/1918.

[37] John Lowengrub, Hermann B. Frieboes, Fang Jin, Yao-Li Chuang, Xiangrong Li, Paul Macklin, Steven M. Wise, and Vittorio Cristini. Nonlinear modeling of cancer: Bridging the gap between cells and tumors. Nonlinearity, 23(1):R1–R91, 2010. doi: 10.1088/0951-7715/23/1/R01. URL http://dx.doi.org/10.1088/0951-7715/23/1/R01. (invited author: J. Lowengrub).

[38] Jonathan Ozik, Nicholson Collier, Randy Heiland, Gary An, and Paul Macklin. Learning-accelerated Discovery of Immune-Tumour Interactions. Molec. Syst. Design Eng., 4:747–60, 2019. doi: 10.1039/c9me00036d. URL http://dx.doi.org/10.1039/c9me00036d.

[39] Ahmadreza Ghaffarizadeh, Samuel H. Friedman, and Paul Macklin. BioFVM: an efficient, parallelized diffusive transport solver for 3-D biological simulations. Bioinformatics, 32(8):1256–8, 2016. doi: 10.1093/bioinformatics/btv730. URL http://dx.doi.org/10.1093/bioinformatics/btv730.

[40] Ahmadreza Ghaffarizadeh, Randy Heiland, Samuel H Friedman, Shannon M Mumenthaler, and Paul Macklin. PhysiCell: an open source physics-based cell simulator for 3-D multicellular systems. PLoS Comput. Biol., 14(2):e1005991, 2018. doi: 10.1371/journal.pcbi.1005991. URL http://dx.doi.org/10.1371/journal.pcbi.1005991.

[41] John Metzcar, Yafei Wang, Randy Heiland, and Paul Macklin. A review of cell-based computational modeling in cancer biology. JCO Clinical Cancer Informatics, 3:1–13, 2019. doi: 10.1200/CCI.18.00069. URL http://dx.doi.org/10.1200/CCI.18.00069. (invited review).

[42] Paul Macklin. PhysiCell demo: immune cells attacking a heterogeneous tumor, visited 2019-11-04. URL https://www.youtube.com/watch?v=nJ2urSm4ilU.

[43] Paul Macklin and Randy Heiland. PhysiCell cancer-immune model (Version 1.3). 2019. doi: 10.21981/M8PR-EC68. URL https://nanohub.org/tools/pc4cancerimmune.

[44] Paul Macklin and Randy Heiland. PhysiCell: invader-scout-attacker system (Version 1.3). 2019. doi: 10.21981/VYDD-NG35. URL https://www.nanohub.org/tools/pcisa.

[45] Gaelle Letort, Arnau Montagud, Gautier Stoll, Randy Heiland, Emmanuel Barillot, Paul Macklin, Andrei Zinovyev, and Laurence Calzone. PhysiBoSS: a multi-scale agent based modelling framework integrating physical dimension and cell signalling. Bioinformatics, 35(7):1188–96, 2019. doi: 10.1093/bioinformatics/bty766. URL http://dx.doi.org/10.1093/bioinformatics/bty766.

[46] Samuel H Friedman, Alexander R A Anderson, David M Bortz, Alexander G Fletcher, Hermann B Frieboes, Ahmadreza Ghaffarizadeh, David Robert Grimes, Andrea Hawkins-Daarud, Stefan Hoehme, Edwin F Juarez, Carl Kesselman, Roeland M H Merks, Shannon M Mumenthaler, Paul K Newton, Kerri-Ann Norton, Rishi Rawat, Russell C Rockne, Daniel Ruderman, Jacob Scott, Suzanne S Sindi, Jessica L Sparks, Kristin Swanson, David B Agus, and Paul Macklin. MultiCellDS: a community-developed standard for curating microenvironment-dependent multicellular data. bioRxiv, 090456, 2016. doi: 10.1101/090456. URL http://dx.doi.org/10.1101/090456. (Preprint, preparing for resubmission).

[47] Paul Macklin. Progress Towards an Open Source PhysiCell Ecosystem, 2019. URL https://dx.doi.org/10.6084/m9.figshare.9934469.v1. October 2, 2019 seminar talk for the Center for Reproducible Biomedical Modeling.

[48] A Madamanchi, Randy Heiland, Paul Macklin, and A J Magana. Students’ Use of Metacognitive Skills in Undergraduate Research Experiences in Computational Modeling. Proceedings–Frontiers in Education Conference, FIE, October 2019, Covington, KY, USA, pages 1–5, 2019. doi: 10.1109/FIE43999.2019.9028386.

[49] Randy Heiland and Paul Macklin. pc4nanobio: cancer nanotherapy simulator (Version 0.8.1). 2018. doi: 10.4231/D3QN5ZD7F. URL https://www.nanohub.org/tools/pc4nanobio.

[50] Randy Heiland, Daniel Mishler, Tyler Zhang, Eric Bower, and Paul Macklin. xml2jupyter: Mapping parameters between XML and Jupyter widgets. Journal of Open Source Software, 4(39):1408, 2019. doi: 10.21105/joss.01408. URL http://dx.doi.org/10.21105/joss.01408.

[51] Gautier Stoll, Barthélémy Caron, Eric Viara, Aurélien Dugourd, Andrei Zinovyev, Aurélien Naldi, Guido Kroemer, Emmanuel Barillot, and Laurence Calzone. MaBoSS 2.0: an environment for stochastic Boolean modeling. Bioinformatics, 33(14):2226–2228, 03 2017. ISSN 1367-4803. doi: 10.1093/bioinformatics/btx123. URL https://doi.org/10.1093/bioinformatics/btx123.

[52] Patrick G Wall. python-loader: Python data loader for PhysiCell digital snapshots, visited 2019-11-04. URL https://github.com/PhysiCell-Tools/Python-loader.

[53] Patrick G Wall. PhysiCell Tools : python-loader (Tutorial Blog Post), visited 2019-11-04. URL http://www.mathcancer.org/blog/python-loader/.

[54] Susan H Russell, Mary P Hancock, and James McCullough. Benefits of undergraduate research experiences. Science, 316(5824):548–549, 2007. doi: 10.1126/science.1140384. URL https://dx.doi.org/10.1126/science.1140384.

[55] Omolola A Adedokun, Ann B Bessenbacher, Loran C Parker, Lisa L Kirkham, and Wilella D Burgess. Research skills and stem undergraduate research students’ aspirations for research careers: Mediating effects of research self-efficacy. Journal of Research in Science teaching, 50(8):940–951, 2013. doi: 10.1002/tea.21102. URL http://dx.doi.org/10.1002/tea.21102.

[56] Wei Zhan. Research experience for undergraduate students and its impact on stem education. Journal of STEM Education, 15(1), 2014. URL https://www.learntechlib.org/p/148288/.

[57] Erin L Dolan and Deborah Johnson. The undergraduate– postgraduate–faculty triad: unique functions and tensions associated with undergraduate research experiences at research universities. CBE—Life Sciences Education, 9(4):543–553, 2010. doi: 10.1187/cbe.10-03-0052. URL https://doi.org/10.1187/cbe.10-03-0052.

[58] Zakiya S Wilson, Lakenya Holmes, Karin Degravelles, Monica R Sylvain, Lisa Batiste, Misty Johnson, Saundra Y McGuire, Su Seng Pang, and Isiah M Warner. Hierarchical mentoring: A transformative strategy for improving diversity and retention in undergraduate stem disciplines. Journal of Science Education and Technology, 21(1):148–156, 2012. doi: 10.1007/s10956-011-9292-5. URL https://dx.doi.org/10.1007/s10956-011-9292-5.

[59] Benjamin Ahn and Monica F Cox. Knowledge, skills, and attributes of graduate student and postdoctoral mentors in undergraduate research settings. Journal of Engineering Education, 105(4):605–629, 2016. doi: 10.1002/jee.20129. URL https://doi.org/10.1002/jee.20129.

[60] Heather Thiry and Sandra L Laursen. The role of student-advisor interactions in apprenticing undergraduate researchers into a scientific community of practice. Journal of Science Education and Technology, 20(6):771–784, 2011. doi: 10.1007/s10956-010-9271-2. URL https://dx.doi.org/10.1007/s10956-010-9271-2.

[61] Cianán B Russell and Gabriela C Weaver. A comparative study of traditional, inquiry-based, and research-based laboratory curricula: impacts on understanding of the nature of science. Chemistry Education Research and Practice, 12(1):57–67, 2011. doi: 10.1039/C1RP90008K. URL https://dx.doi.org/10.1039/C1RP90008K.

[62] Elaine Seymour, Anne-Barrie Hunter, Sandra L Laursen, and Tracee DeAntoni. Establishing the benefits of research experiences for undergraduates in the sciences: First findings from a three-year study. Science education, 88(4):493–534, 2004. doi: 10.1002/sce.10131. URL https://dx.doi.org/10.1002/sce.10131.

[63] Jenny Olin Shanahan, Elizabeth Ackley-Holbrook, Eric Hall, Kearsley Stewart, and Helen Walkington. Ten salient practices of undergraduate research mentors: A review of the literature. Mentoring & Tutoring: Partnership in Learning, 23(5):359–376, 2015. doi: 10.1080/13611267.2015.1126162. URL https://dx.doi.org/10.1080/13611267.2015.1126162.

[64] D Brian Larkins, J Christopher Moore, Louis J Rubbo, and Laura R Covington. Application of the cognitive apprenticeship framework to a middle school robotics camp. In Proceeding of the 44th ACM technical symposium on Computer science education, pages 89–94. ACM, 2013. doi: 10.1145/2445196.2445226. URL https://dx.doi.org/10.1145/2445196.2445226.

[65] Susan Howitt and Anna Wilson. Scaffolded reflection as a tool for surfacing complex learning in undergraduate research projects. Council on Undergraduate Research Quarterly, 36(4), 2016. doi: 10.18833/curq/36/4/8.

[66] Mark Guzdial. Software-realized scaffolding to facilitate programming for science learning. Interactive learning environments, 4.1:1–44, 1994. URL http://guzdial.cc.gatech.edu/Emile-ILE.pdf.

[67] Hayden W Fennell, Joseph A Lyon, Aasakiran Madamanchi, and Alejandra J Magana. Toward computational apprenticeship: Bringing a constructivist agenda to computational pedagogy. Journal of Engineering Education, 109(2):170–176, 2020. doi: 10.1002/jee.20316.

[68] Sam Donovan, Carrie Diaz Eaton, Stith T. Gower, Kristin P. Jenkins, M. Drew LaMar, DorothyBelle Poli, Robert Sheehy, and Jeremy M. Wojdak. QUBES: a community focused on supporting teaching and learning in quantitative biology. Letters in Biomathematics, 2(1):46–55, 2015. doi: 10.1080/23737867.2015.1049969. URL https://doi.org/10.1080/23737867.2015.1049969.

[69] Paul Macklin and Randy Heiland. PhysiCell: biased random migration demonstrator (Version 1.1). 2019. doi: 10.21981/AW4Z-VN50. URL https://www.nanohub.org/tools/motility.

[70] Furkan Kurtoglu, Aneequa Sundus, Kali Nicole Konstantinopoulos, Drew Willis, Mary Chen, Randy Heiland, and Paul Macklin. Motility Training App for PhysiCell (Version 1.1). 2019. doi: 10.21981/RQWR-T970. URL https://www.nanohub.org/tools/trmotility.

